# Megakaryocyte emperipolesis mediates membrane transfer from intracytoplasmic neutrophils to platelets

**DOI:** 10.1101/504555

**Authors:** Pierre Cunin, Rim Bouslama, Kellie R. Machlus, Marta Martínez-Bonet, Pui Y. Lee, Nathan Nelson-Maney, Allyn Morris, Li Guo, Andrew S. Weyrich, Martha Sola-Visner, Eric Boilard, Joseph E. Italiano, Peter A. Nigrovic

**Author notes:** Correspondence – Lead contact, Peter A. Nigrovic, MD, Associate Professor of Medicine, Harvard Medical School, Brigham and Women’s Hospital, 60 Fenwood Road, BTM6002L, Boston MA 02115 USA, www.nigroviclab.org.

## Abstract

Bone marrow megakaryocytes engulf neutrophils in a phenomenon termed emperipolesis. We show here that emperipolesis is a dynamic process mediated actively by both lineages, in part through the β2-integrin/ICAM-1/ezrin pathway. Tethered neutrophils enter in membrane-bound vesicles before penetrating into the megakaryocyte cytoplasm. Intracytoplasmic neutrophils develop membrane contiguity with the demarcation membrane system, thereby transferring membrane to the megakaryocyte and to daughter platelets. This phenomenon occurs in otherwise unmanipulated marrow *in vivo*, resulting in circulating platelets that bear membrane from non-megakaryocytic hematopoietic donors. Transit through megakaryocytes can be completed as rapidly as minutes, after which neutrophils egress intact. Emperipolesis is amplified in models of inflammation associated with platelet overproduction, contributing to platelet production *in vitro* and *in vivo.* These findings identify emperipolesis as a new cell-in-cell interaction that enables neutrophils and potentially other cells passing through the megakaryocyte cytoplasm to modulate the production and membrane content of platelets.

## Introduction

Megakaryocytes (MKs) are the cellular source of platelets. Derived from hematopoietic stem cells, developing MKs undergo multiple rounds of endomitosis to become highly-polyploid cells averaging 20 to 100 μm in size (Levine et al., 1982; Machlus and Italiano, 2013). Mature MKs develop a complex network of intracytoplasmic membrane, termed the demarcation membrane system (DMS), that provides a membrane reservoir to enable platelet generation (Schulze et al., 2006). MKs then protrude pseudopodial extensions of this membrane via the marrow sinusoids into the bloodstream, where shear stress releases fragments that become the mature platelets required for hemostasis (Junt et al., 2007).

Representing less than 0.3% of hematopoietic cells in bone marrow, (Levine et al., 1982; Machlus and Italiano, 2013; Winter et al., 2010) MKs interact with other hematopoietic lineages. MKs provide a niche for plasma cells (Winter et al., 2010), promote neutrophil egress via production of CXCR2 ligand (Kohler et al., 2011), and regulate hematopoietic stem cell homeostasis (Bruns et al., 2014; Zhao et al., 2014). Almost 50 years ago, it was observed that MKs can engulf other hematopoietic cells, most commonly neutrophils (Larsen, 1970). Examination of fresh aspirates revealed movement of these cells within MKs, giving rise to the name emperipolesis from the Greek, *em* inside, *peri* around, *polemai* wander about (Humble et al., 1956; Larsen, 1970) Emperipolesis is observed in healthy marrow and increases with hematopoietic stress, including myelodysplastic and myeloproliferative disorders (Cashell and Buss, 1992; Mangi and Mufti, 1992), myelofibrosis (Centurione et al., 2004; Schmitt et al., 2002; Spangrude et al., 2016), gray platelet syndrome (Di Buduo et al., 2016; Larocca et al., 2015; Monteferrario et al., 2014), essential thrombocythemia (Cashell and Buss, 1992), blood loss or hemorrhagic shock (Dzieciol et al., 1995; Sahebekhitiari and Tavassoli, 1976; Tavassoli, 1986). Its mechanism and significance remain unknown. It has been speculated that MKs could represent a sanctuary for neutrophils in an unfavorable marrow environment, or a route for neutrophils to exit the bone marrow, but more typically emperipolesis is regarded as a histological curiosity without physiological significance (Lee, 1989; Sahebekhitiari and Tavassoli, 1976; Tavassoli, 1986).

Recently, we identified evidence for a direct role for MKs in systemic inflammation, highlighting the potential importance of the interaction of MKs with immune lineages (Cunin et al., 2017). Whereas the preservation of emperipolesis in monkeys (Stahl et al., 1991), mice (Centurione et al., 2004), rats (Tanaka et al., 1996), and cats and dogs (Scott and Friedrichs, 2009) suggests evolutionary conservation, we sought to model this process *in vitro* and *in vivo* to begin to understand its biology and function. We show here that emperipolesis is a tightly-regulated process mediated actively by both MKs and neutrophils via pathways reminiscent of leukocyte transendothelial migration. Neutrophils enter MKs within membrane-bound vesicles but then penetrate into the cell cytoplasm, where they develop membrane continuity with the DMS to transfer membrane to MKs and thereby to platelets, a process that accelerates platelet production. Neutrophils then emerge intact, carrying MK components with them. Together, these data identify emperipolesis as a previously unrecognized form of cell-in-cell interaction that mediates a novel mode of material transfer between immune and hematopoietic lineages.

## Results

### *In vitro* modeling of emperipolesis reveals a rapid multi-stage process

Whole-mount 3D immunofluorescence imaging of healthy C57Bl/6 murine marrow revealed that ~6% of MKs contain at least one neutrophil, and occasionally other bone marrow cells (Fig 1A and Movie 1). Emperipolesis was similarly evident upon confocal imaging of unmanipulated human marrow (Fig 1B). To model this process, we incubated cultured murine or human MKs with fresh bone marrow cells or peripheral blood neutrophils, respectively (Fig 1C&D). Murine MKs, derived either from bone marrow or fetal liver cells, were efficient at emperipolesis (~20-40% of MKs). Neutrophils were by far the most common participants, although B220+ B cells, CD115+ monocytes, and occasional CD3+ T cells and NK1.1+ NK cells were also observed within MKs (Fig S1A). Emperipolesis was less efficient in human cultured MKs (2-5% of MKs), which are typically much smaller than murine MKs, and was observed in MKs cultured from marrow CD34+ cells but not from the even smaller MKs derived from cord blood CD34+ cells (Fig 1D and not shown). We elected to continue our mechanistic studies in murine MKs, principally cultured from marrow.

**Figure 1:**
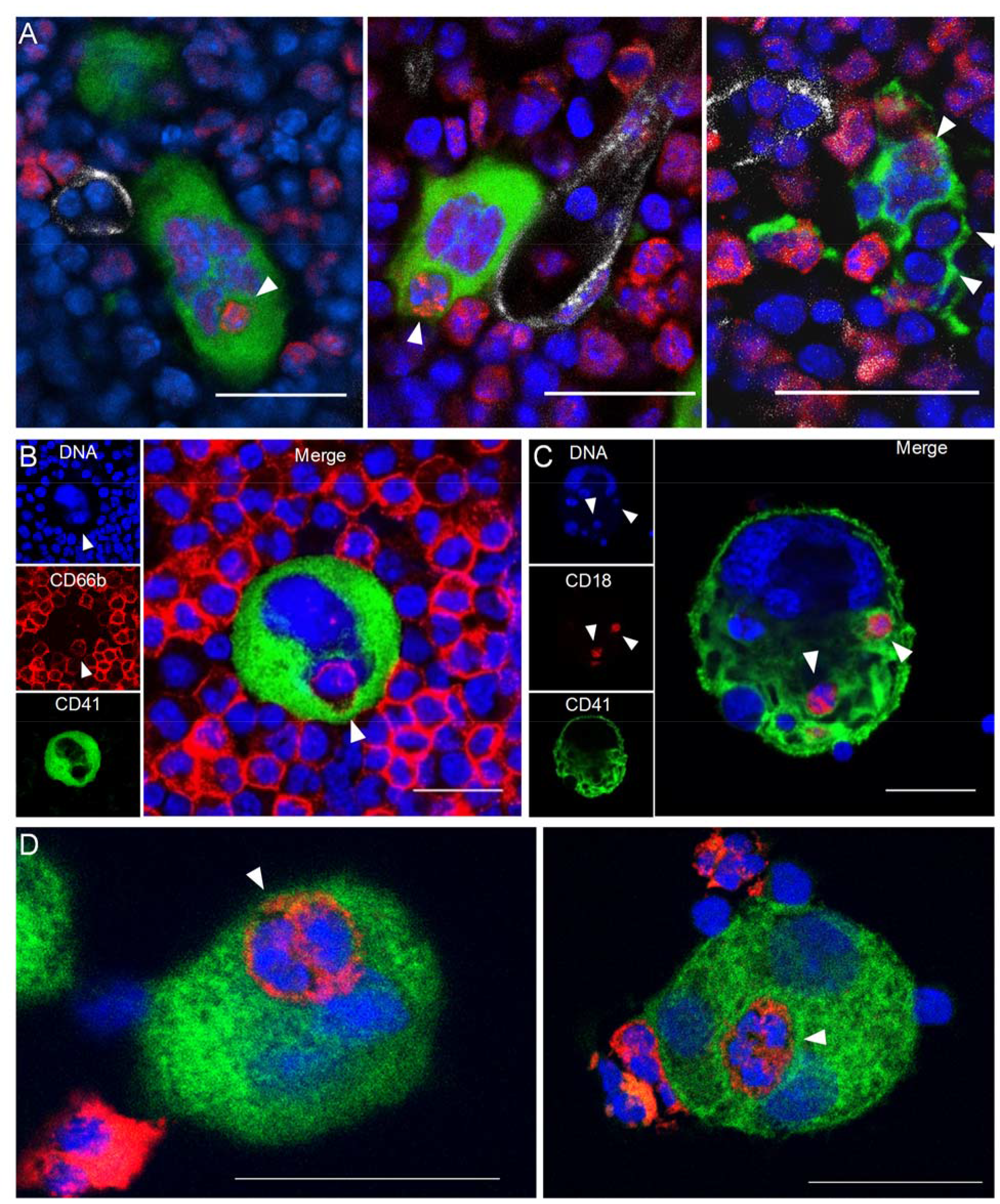
Visualization of murine and human emperipolesis by confocal microscopy. **A**. Whole-mount images of mouse bone marrow stained with anti-CD41 (green), anti-Ly6G (red) and anti-CD31/CD144 (white). Arrowheads show internalized neutrophils or other Ly6G^neg^ bone marrow cells (right image). Three-dimensional reconstitutions and confirmation of cell internalization are shown in Supplemental movie 1. **B**. Cells from human bone marrow aspirate were stained with anti-CD41 (green) and anti-CD66b (red). **C**. Murine MKs were co-cultured with marrow cells overnight. Cells were stained with anti-CD41 (green) and anti-CD18 (red). **D**. Human MKs generated from marrow CD34+ cells were co-cultured with circulating neutrophils overnight. Cells were stained with anti-CD41 (green) and anti-CD15 (red). **A-D**. DNA was visualized with Draq5 or Hoechst (blue), arrowheads represent internalized neutrophils, scale bars represent 20μm.

### Neutrophils engaged in emperipolesis penetrate into the MK cytoplasm

Confocal microscopy revealed four distinct steps. First, neutrophils become adherent to the MK surface (Fig 2A and Movie 2&3), including to membrane protrusions we term *MK tethers* (Fig S1B and movie 3). Second, neutrophils enter MKs within membrane-bound vacuoles, hereafter termed *emperisomes*, bearing the MK surface marker CD41+ (Fig 2A and Movie 3). Third, the emperisome undergoes transformation such that CD41 is no longer evident (Fig 2A and Movie 4). While most MKs engaged in emperipolesis contained only 1 or 2 neutrophils, some resembled “reservoirs” containing dozens of neutrophils in stages 2 and 3 (Fig S1C), an appearance recognized in human marrow as well (Cashell and Buss, 1992; Larsen, 1970; Monteferrario et al., 2014; Thiele et al., 1984) (Fig S1D). Fourth, neutrophils exited MKs, returning to the extracellular milieu as viable motile cells (Fig 2A, Fig S4D and Movies 3-5). Live cell imaging of murine MKs co-incubated with fresh bone marrow cells showed that neutrophil transit was of variable duration, in some cases lasting only a few minutes (Fig S1E, Fig S4D and Movie 2, 3&5) and in others more than one hour (Movie 5&6).

**Figure 2:**
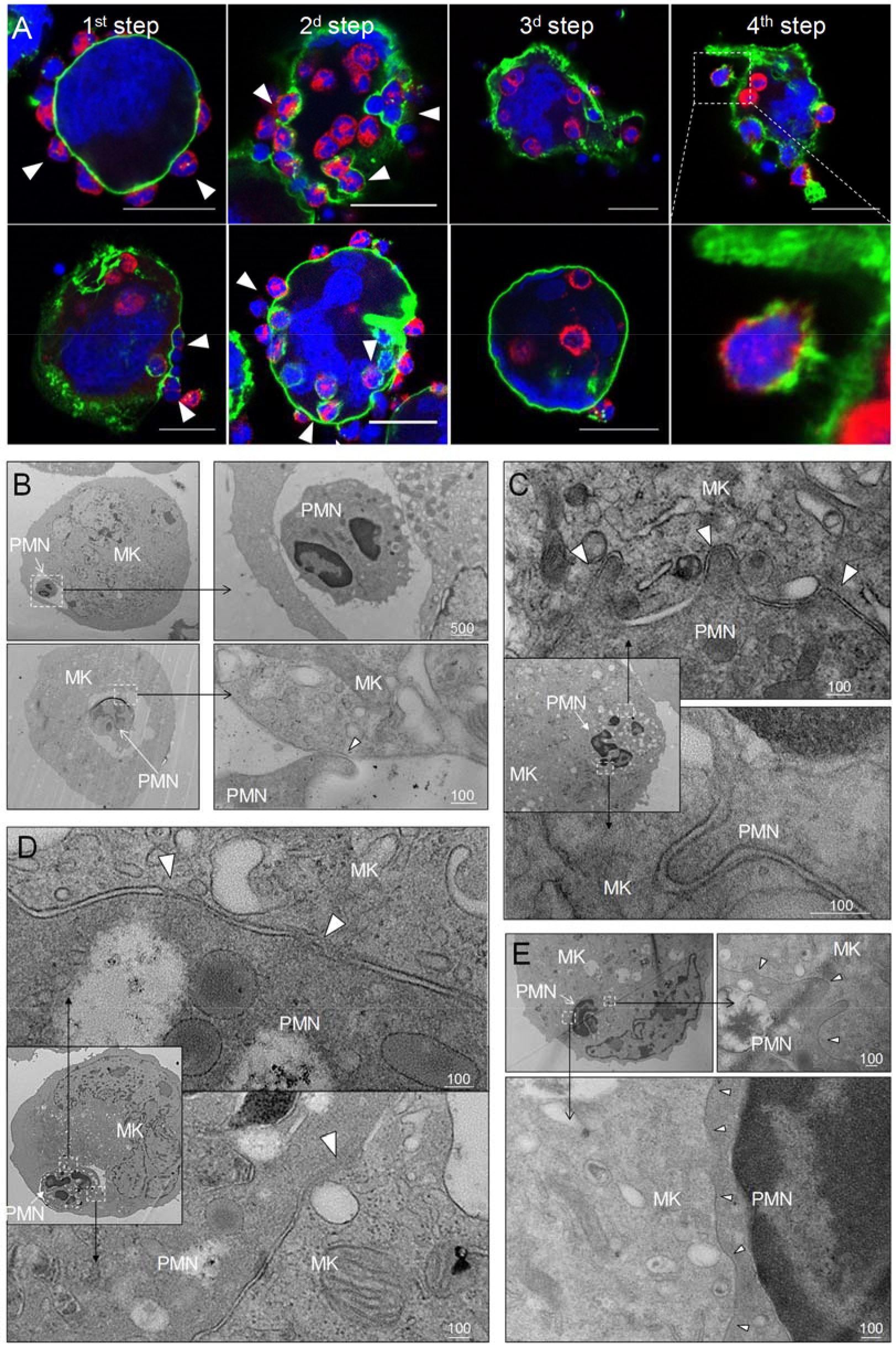
In vitro modeling of emperipolesis reveals a rapid multi-stage process. **A**. Confocal images showing different steps of emperipolesis. Scale bars represent 20μm. **B-E**. Cells were stained with OsO4 after emperipolesis assay for electron microscopy observation. B. Neutrophils inside MK vacuoles. Some neutrophil surface protrusions make contact with the emperisome membrane (arrowhead). **C-D**. Close interaction between neutrophil and emperisome membrane forming a two-membrane leaflet structure. **C**. Protrusions from neutrophils that extend inside the MK cytoplasm (arrowheads). **D**. Demarcation between emperisome membrane and neutrophil membrane disappears (arrowheads). **E**. Neutrophil and MK are separated by a single membrane (arrowheads). **B-E**. Scale bars in nm.

To better understand the stages of emperipolesis, we employed electron microscopy (EM). After neutrophil uptake into the emperisome (Fig 2B), the vacuolar space between neutrophil and MK was resorbed such that neutrophil and MK membranes became closely apposed, resulting in a structure composed of two membrane leaflets surrounding the neutrophil (Fig 2C&D). This structure was often associated with the appearance of neutrophilic protrusions deeper into the host MK (Fig 2C). Areas in which the membranes approximated very closely, becoming indistinct for short stretches, were sometimes observed (Fig 2D). Subsequently, only a single bilipid membrane came to separate the neutrophil cytoplasm from the MK cytoplasm, a finding that echoed the loss of CD41 staining observed by immunofluorescence, confirming dissolution of the emperisome and thereby translocation of the neutrophil to an intracytoplasmic location (Fig 2E). Whereas CD18 and Ly6G but not CD41 were preserved (Fig 2A above), this remaining membrane is most likely primarily of neutrophil origin.

### Emperipolesis is mediated by active actin cytoskeleton rearrangement in both megakaryocyte and neutrophil

To assess the cytoskeletal processes underlying this intriguing cell-in-cell interaction, we employed targeted inhibitors. The microtubule polymerization inhibitor nocodazole showed a negligible effect, but emperipolesis was dramatically curtailed by inhibitors of actin polymerization, cytochalasin D and latrunculin A (Fig 3A). This effect was observed when either MKs or marrow cells were exposed to these inhibitors, confirming obligate active cytoskeletal engagement by both participants (Fig 3B, controls of actin inhibition in Fig S2A). Consistent with these results, neutrophils entering MKs exhibited a polarized appearance, while MKs developed a transcellular cup similar to that observed in endothelial cells during transendothelial migration of leukocytes (Carman and Springer, 2004; Ley et al., 2007) (Fig S1E, Fig S2B, and Movie 2). In agreement with inhibitor findings, actin but not microtubules localized to the interface between MKs and extracellular neutrophils, and was observed to encase neutrophil-containing CD41+ vacuoles (Fig 3D and Fig S2C); by contrast, actin was not observed surrounding neutrophils that were no longer delimited by CD41+ membrane (Fig 3C). These observations demonstrate that emperipolesis is an active process mediated by actin cytoskeletal rearrangement of both MKs and neutrophils.

**Figure 3:**
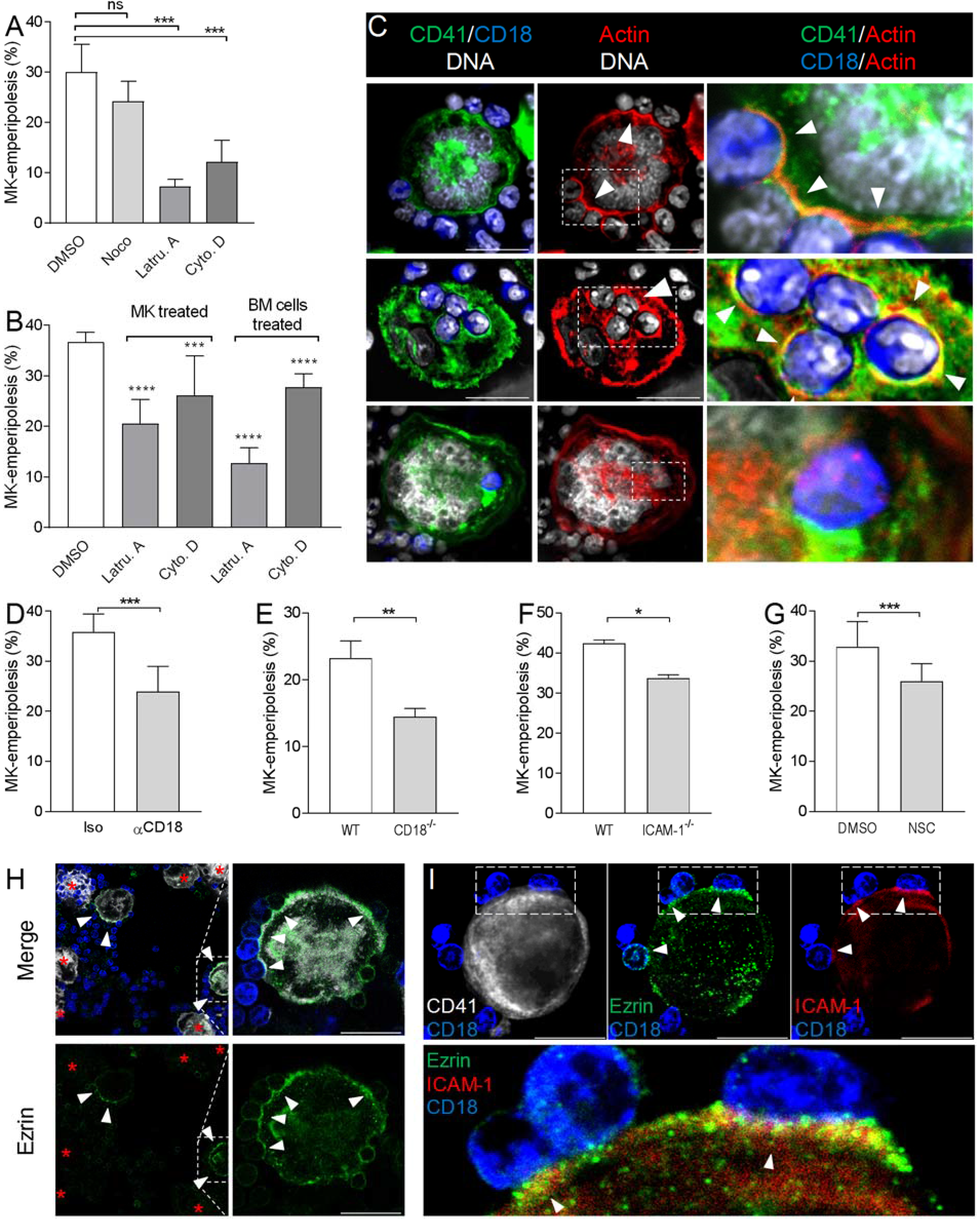
Emperipolesis is mediated by active actin cytoskeleton rearrangement and in part through β2-integrin/ICAM-1/ezrin. **A**. MK and marrow cells were co-cultured in the presence of 1μM nocodazole, latrunculin A, cytochalasin D, or control vehicle. **B**. MKs or marrow cells were treated with latrunculin A or cytochalasin D for two hours. After thorough washings, cells were co-cultured with untreated marrow cells or MKs, respectively. **A-B**. Cells are stained with anti-CD41, -CD18 or -Ly6G and Draq5, and observed by confocal microscopy. Histograms show percentages of MKs containing at least one neutrophil. At least 150 (**A**) or 500 (**B**) MKs per condition were counted; pool of 3 independent experiments. **C**. Cells are stained with anti-CD41 (green), anti-CD18 (blue) and phalloidin (red). DNA is visualized with Hoechst (gray). Images show F-actin on MK surface where neutrophils are attached (upper photos), around neutrophils encapsulated in CD41+ vacuoles (middle photos) or free within MKs (lower photos). **D-G**. Emperipolesis assay was performed (**A**) in the presence of 10μg/ml anti-CD18 or corresponding isotype control rat IgG1 (**B**) using marrow cells from WT versus CD18-deficient mice or (**C**) using MKs from WT versus ICAM-1-deficient mice, or (**D**) in the presence of 1μM of ezrin inhibitor NSC668394. **A-D**. Histograms show percentages of MKs containing at least one neutrophil. At least 350 MKs per condition were counted; pool of 2 (**F**), 3 (**E**), or 4 (**D&G**) independent experiments. **H**. After co-culture, cells are stained with anti-CD41 (white), anti-ezrin (green) and anti-Ly6G (blue). Arrows show ezrin clustering on the surface of MK. Red asterisk show MK without detectable ezrin. **I**. Cells are stained with anti-CD41 (gray), -ezrin (green), – ICAM-1 (red), -Ly6G (blue). Arrows show ICAM-1/ezrin co-localization on MK surface. Lower photo is a magnification of the dashed white region. **H&I**. Scale bars represent 20μm.

### Emperipolesis is mediated in part through β2-integrin/ICAM-1/ezrin

To define how MKs and neutrophils interact, we tested several candidate ligand/receptors pathways. Blocking antibodies targeting P-selectin, glycoprotein VI (GPVI), PECAM-1, CD44, and CXCR2 had no effect (not shown). However, blockade of the β2 integrin CD18, expressed by neutrophils but not MKs, strongly impaired emperipolesis (Fig 3D). Correspondingly, CD18-deficient bone marrow cells exhibited reduced emperipolesis into WT MKs (Fig 3E), despite the heightened proportion and density of neutrophils in these marrows (Horwitz et al., 2001).

β2 integrins bind ICAM-1, among other targets (Ley et al., 2007). Confocal microscopy showed that ICAM-1 is expressed by a population of human and murine MKs (Fig S3A-D). In agreement with previous observations in rat (Tanaka et al., 1997), emperipolesis by ICAM-1-deficient MKs was significantly impaired (Fig 3F). In further support of this mechanism, we evaluated the role of ezrin, which mediates the attachment of the intracellular tail of ICAM-1 to the actin cytoskeleton (Heiska et al., 1998; Ley et al., 2007). Inhibition of ezrin phosphorylation impaired emperipolesis (Fig 3G, controls of ezrin inhibition in Fig S3E). By confocal microscopy, ezrin could be detected only at sites of MK contact with tethered neutrophils, where it co-localized strongly with ICAM-1, consistent with its role as a bridge to the cytoskeleton (Fig 3H&I). By contrast, ezrin could not be visualized in MKs simply tethered to leukocytes, or with only internalized leukocytes (Fig 3H). Together, these data show that emperipolesis is mediated in part by an interaction between neutrophil β2 integrins and MK ICAM-1/ezrin during neutrophil entry. However, absence or blockade of these factors resulted in only partial impairment of emperipolesis, indicating a role for alternate mechanisms not yet defined.

### Emperipolesis mediates membrane transfer from neutrophil to megakaryocyte

As observed by others (Centurione et al., 2004; Thiele et al., 1984), neutrophils engaged in emperipolesis frequently localized to the DMS, the intracytoplasmic membrane network implicated in platelet production (Fig 4A). Close examination of this interaction demonstrated membrane contiguity between the neutrophil and the DMS, suggesting that neutrophils could serve as membrane donors to MKs and potentially to platelets (Fig 4A). To test this possibility, we employed membrane labeling. Marrow cells were stained with the lipophilic dye CellVue maroon and then co-cultured with unstained MKs. Confocal microscopy showed substantial loss of CellVue maroon staining in neutrophils inside but not outside MKs (Fig 4B). Similar loss of fluorescence was observed with the lipid stains Bodipy and PKH67, excluding a non-specific chemical interaction (Fig S4A&B). Further, some cells exhibited diffusion of stain within the MK (Fig 4B & S4C), consistent with the membrane transfer implied by EM. Finally, time-lapse spinning disk confocal microscopy confirmed transfer of neutrophil membrane to MKs from this intracellular location (Fig S4D and Movies 5&6).

**Figure 4:**
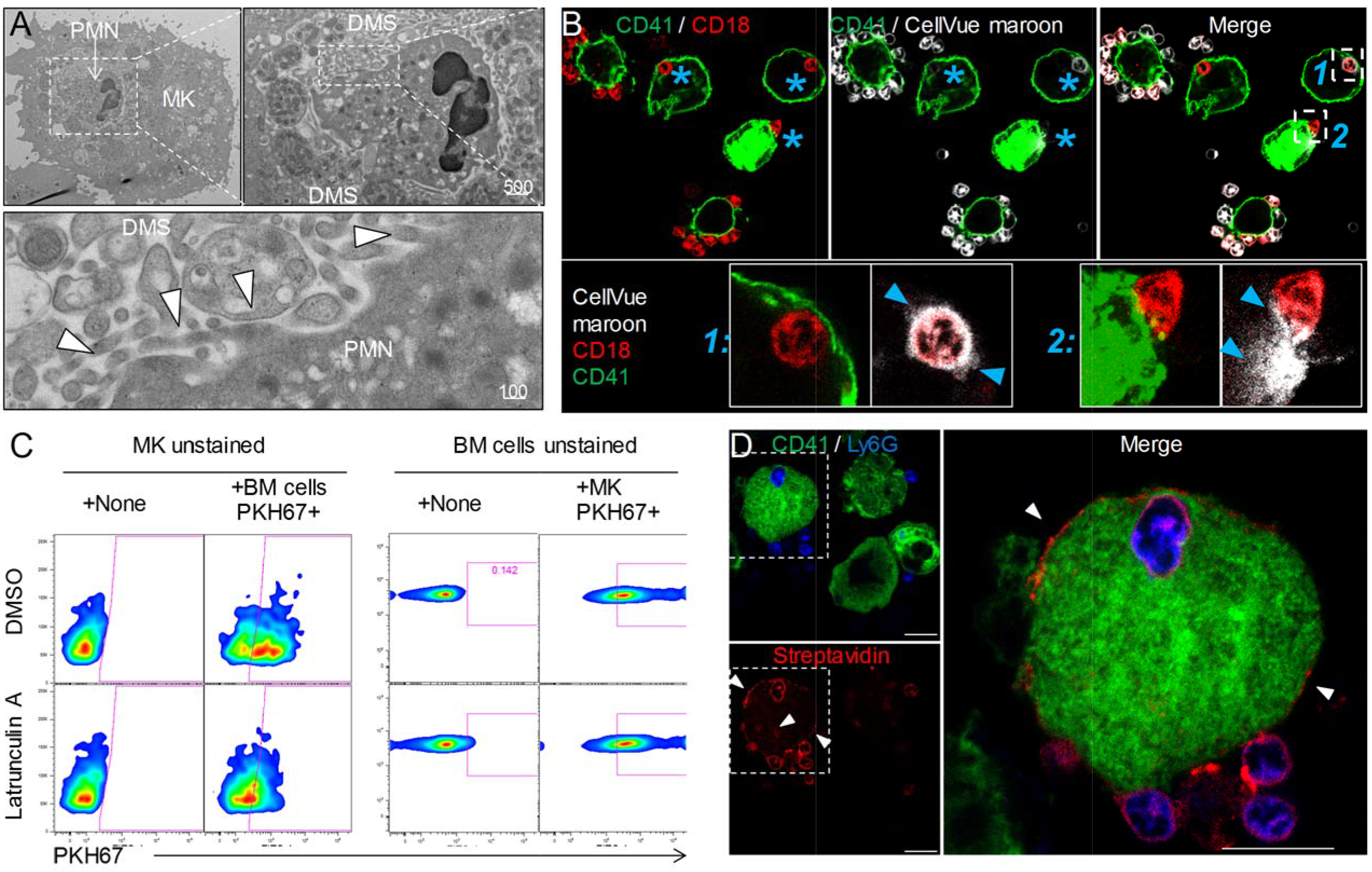
Emperipolesis mediates membrane transfer from neutrophil to megakaryocyte. **A**. Cells were stained with OsO4 after emperipolesis assay for electron microscopy observation. Images show a membrane continuity between neutrophil and DMS (arrowheads). **B**. Bone marrow cells were stained with CellVue Maroon prior to co-culture with MK. Confocal microscopy shows a loss of CellVue Maroon fluorescence in neutrophils engaged in emperipolesis (blue asterisks). Lower photos show details for the two neutrophils (1) and (2) on the upper right photo (transfer of CellVue Maroon inside MK, blue arrowheads). **C**. MKs and marrow cells were co-cultured with 1μM latrunculin A or DMSO. Left panels: marrow cells are previously stained with PKH67, dot plots show PKH67 fluorescence on CD41+ MKs. Right panels: MKs are previously stained with PKH67, dot plots show PKH67 fluorescence on Ly6G+ neutrophils. **D**. Surface proteins of marrow cells were biotinylated prior to emperipolesis assay. After fixation, cells were incubated with AF594-streptavidin (red). Asterisks show the presence of biotinylated proteins on MK surface and DMS. **B & D**: Scale bars represents 20μm.

To assess reciprocal membrane transfer from MKs to neutrophils, we stained MKs with lipid stains as above and co-cultured these with unstained marrow cells. MK-derived lipids strongly co-localized with neutrophil membrane during emperipolesis (Fig S4E-G), while time-lapse spinning disk microscopy also confirmed reciprocal membrane exchange (Fig S4D and Movie 5&6). While lipid exchange from neutrophils to MKs was strongly inhibited by latrunculin A, transfer from MKs to neutrophils was not (Fig 4C), suggesting that the transfer from MK to neutrophil was not primarily mediated by emperipolesis. MKs produce microparticles in great abundance (Cunin et al., 2017; Flaumenhaft et al., 2009), and PKH67-stained MKs were observed to release many PKH67+ microparticles in a latrunculin A-independent manner that could transfer membrane fluorescence to neutrophils in the absence of intact MKs (Fig S4H). By contrast, no fluorescence was detected on MKs cultured with supernatant from PKH67-stained marrow cells, rendering unlikely a role of marrow cell-derived microparticles, exosomes or apoptotic bodies in membrane transfer from neutrophils to MKs (Fig S3I). Thus, neutrophils transfer plasma membrane to MKs via emperipolesis, while reciprocal membrane exchange from MKs to neutrophils likely occurs primarily via MK microparticles, a phenomenon of interest but not explored further here.

We then sought to determine whether membrane transfer mediates exchange of surface proteins. We performed surface biotinylation of MKs and marrow cells, and then co-cultured these cells with unstained marrow cells or MKs, respectively. Using streptavidin, we could not detect biotin on neutrophils incubated with biotinylated MKs, excluding bulk surface protein transfer from MKs to neutrophils (not shown). However, surface biotin could be detected on some MKs after incubation with biotinylated marrow cells (Fig 4D), confirming that membrane exchange from neutrophils to MK transfers proteins from one cell to the other. The nature of these proteins remains to be determined since MKs remained negative for hallmark neutrophil proteins such as CD18 and Ly6G (not shown).

### Neutrophil membranes transferred in emperipolesis emerge on circulating platelets

Platelets are generated by MKs via the DMS network, an impressively extended network of membrane whose biogenesis remains incompletely understood (Eckly et al., 2014; Schulze et al., 2006). EM had demonstrated membrane continuity between cytoplasmic neutrophils and the DMS. We therefore tested whether emperipolesis could transfer neutrophil membrane to platelets. MKs require shear stress for physiological platelet biogenesis, rendering the *in vivo* context most suitable for these studies. MKs stained with the cytoplasmic dye Green-CMFDA were incubated with marrow cells stained with the lipid marker CellVue Maroon and then engrafted intravenously into congenic recipient mice, in which production of CMFDA+ platelets was monitored by serial phlebotomy (Cunin et al., 2017; Fuentes et al., 2010; Zhang et al., 2016) and Fig S5A). Remarkably, most platelets produced by donor MKs (i.e. CMFDA+) were also positive for CellVue Maroon, indicating a high frequency of incorporation of donor leukocyte membrane (Fig 5A). The intensity of CellVue Maroon staining remained constant over time, suggesting that donor membrane was employed continuously over an extended period (Fig 5B). Similar findings were obtained with lipid stainer PKH67 (Fig S5B). To exclude experimental artifact related to lipid stains, we employed donor marrow from mT/mG mice bearing membrane fluorescence mediated by fluorochrome associated with the inner membrane leaflet. Confocal imaging confirmed that membrane fluorescence from mT/mG marrow cells efficiently transferred into MKs *in vitro* (Fig S5C). Membrane fluorescence was also detected on platelets produced *in vivo* by WT MKs incubated with mT/mG marrow donors, albeit with weaker signal since membrane fluorescence is less intense that with lipid stains (Fig S5D). We similarly investigated transfer of protein. Marrow cells were stained with the intracellular protein stain CellTrace Violet and then co-cultured with CMFDA+ MKs. Interestingly, platelets emerging *in vivo* contained CellTrace violet, consistent with cytoplasmic protein transfer (Fig 5C&D). These results demonstrate that lipids and intracellular proteins are transferred from marrow cells not only to MKs but also to their daughter platelets.

**Figure 5:**
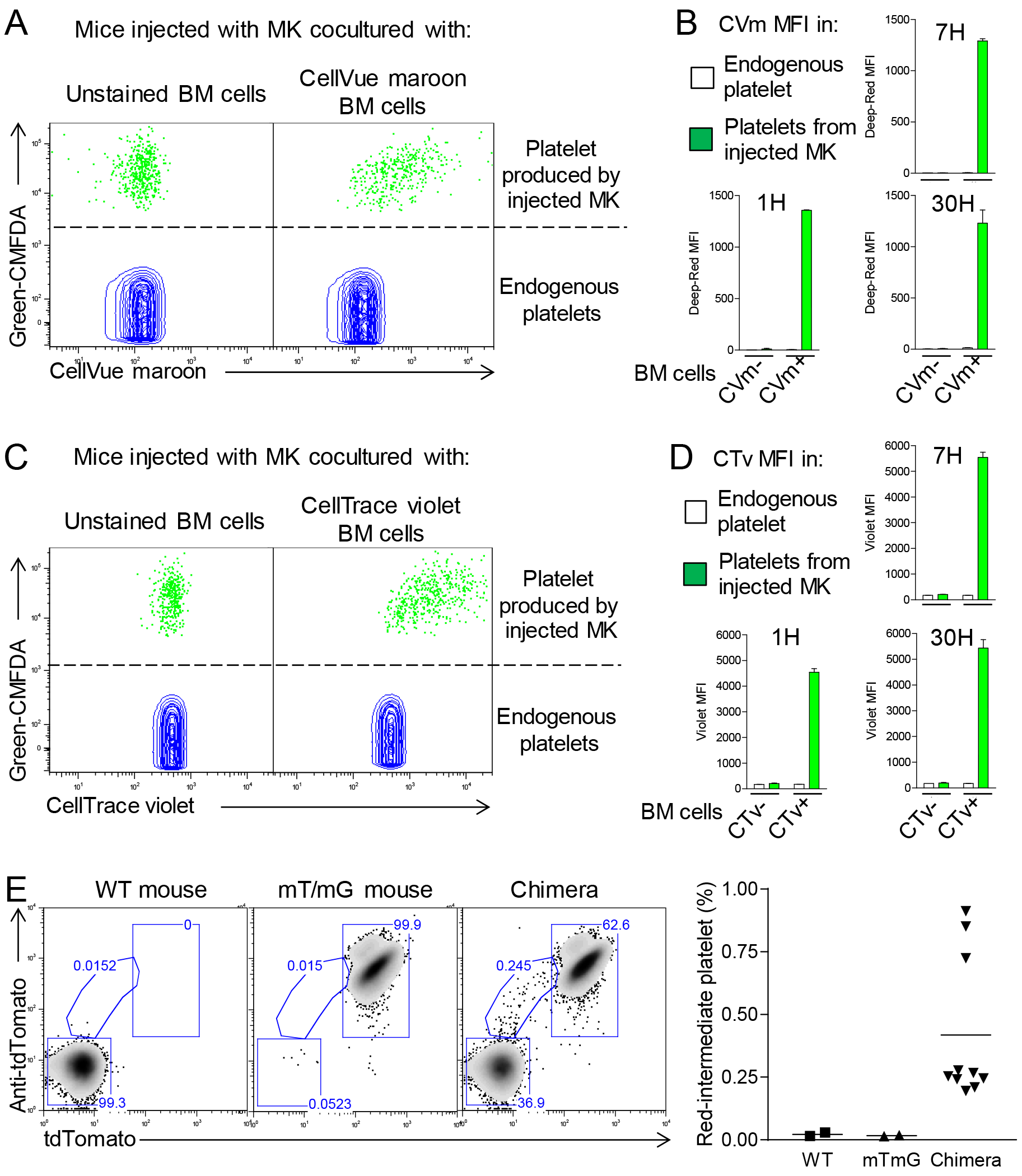
Neutrophil membranes transferred in emperipolesis emerge on circulating platelets. **A**. CMFDA+ MKs are co-cultured with either CellVue maroon (CVm) or unstained marrow cells, and injected i.v. (Figure S5A). Circulating CD41+ platelets were analyzed serially by flow cytometry. Dot plots show CVm MFI among CMFDA-platelets (endogenous) and CMFDA+ platelets (produced by injected MKs). **B**. CellVue maroon MFI on endogenous platelets versus platelets produced by injected MKs over time. **C-D**. Same as (**B-C**) but marrow cells were stained with the protein stain CellTrace violet (CTv) prior to co-culture with CMFDA+ MK. **E**. CD45.1 mice were lethally irradiated and engrafted for 4 weeks with a mix of WT and mT/mG marrow. Circulating platelets were fixed, permeabilized and stained with anti-tdTomato. Left: representative dot plots of tdTomato and anti-tdTomato MFI in circulating platelets from WT, mT/mG, and chimera mice. Right: frequency of the red-intermediate population among total platelets, representative of 3 independent experiments. Note that this population underestimates the true post-emperipolesis population because it detects only mT/mG➔WT events, not WT➔WT, mT/mG➔mT/mG, and WT➔mT/mG.

We sought to exclude the possibility that this membrane transfer reflected an artifact of *ex vivo* MK generation and co-culture. To this end, we generated mice chimeric for WT and mT/mG marrow, allowing us to seek platelets resulting from mT/mG➔WT membrane transfer in a fully native environment. Indeed, platelets with the expected intermediate fluorescent phenotype were observed (Fig 5E), albeit only in relatively small numbers, potentially because of the weak fluorescence in the mT/mG system and because WT➔WT, mT/mG➔mT/mG, and WT➔mT/mG remain undetectable. Examination of BM MKs identified examples of fluorescent neutrophils contributing membrane to non-fluorescent MKs from an intracellular location (Fig S5E). We conclude that intracellular neutrophils transfer membrane to MKs and thereby to platelets via emperipolesis *in vivo.*

### Emperipolesis contributes to platelet production

As a bidirectional interaction between MKs and leukocytes, emperipolesis is likely to have multiple cellular effects. Among these, we elected to explore its impact on thrombocytogenesis. Recognizing the association of emperipolesis in humans with hematopoietic disease (Cashell and Buss, 1992; Centurione et al., 2004; Larsen, 1970; Mangi and Mufti, 1992; Stahl et al., 1991; Thiele et al., 1984), we exposed mice to several models of stress-induced platelet overproduction by MKs, intraperitoneal LPS injection and IgG-mediated thrombocytopenia. The proportion of MKs containing at least one neutrophil was assessed in two-dimensional marrow sections. In each case, emperipolesis increased from a baseline of ~2-5% in control mice to ~6-10% under stress (Fig 6A-C). These figures represent a minimal estimate of the “snapshot” prevalence of emperipolesis, since they sample only 5μm sections of MKs with a typical diameter of 20-100μm, but nevertheless confirm that emperipolesis is common and strongly induced under physiological stress. Interestingly, an enhanced drive for platelet production was not sufficient to augment emperipolesis, because accelerated platelet production following administration of thrombopoietin or platelet depletion by anti-CD41 was unaccompanied by an increase in emperipolesis (Fig S6A&B). One possible explanation is that neutrophil activation may also be required, consistent with the role of neutrophil β2 integrins defined above.

**Figure 6:**
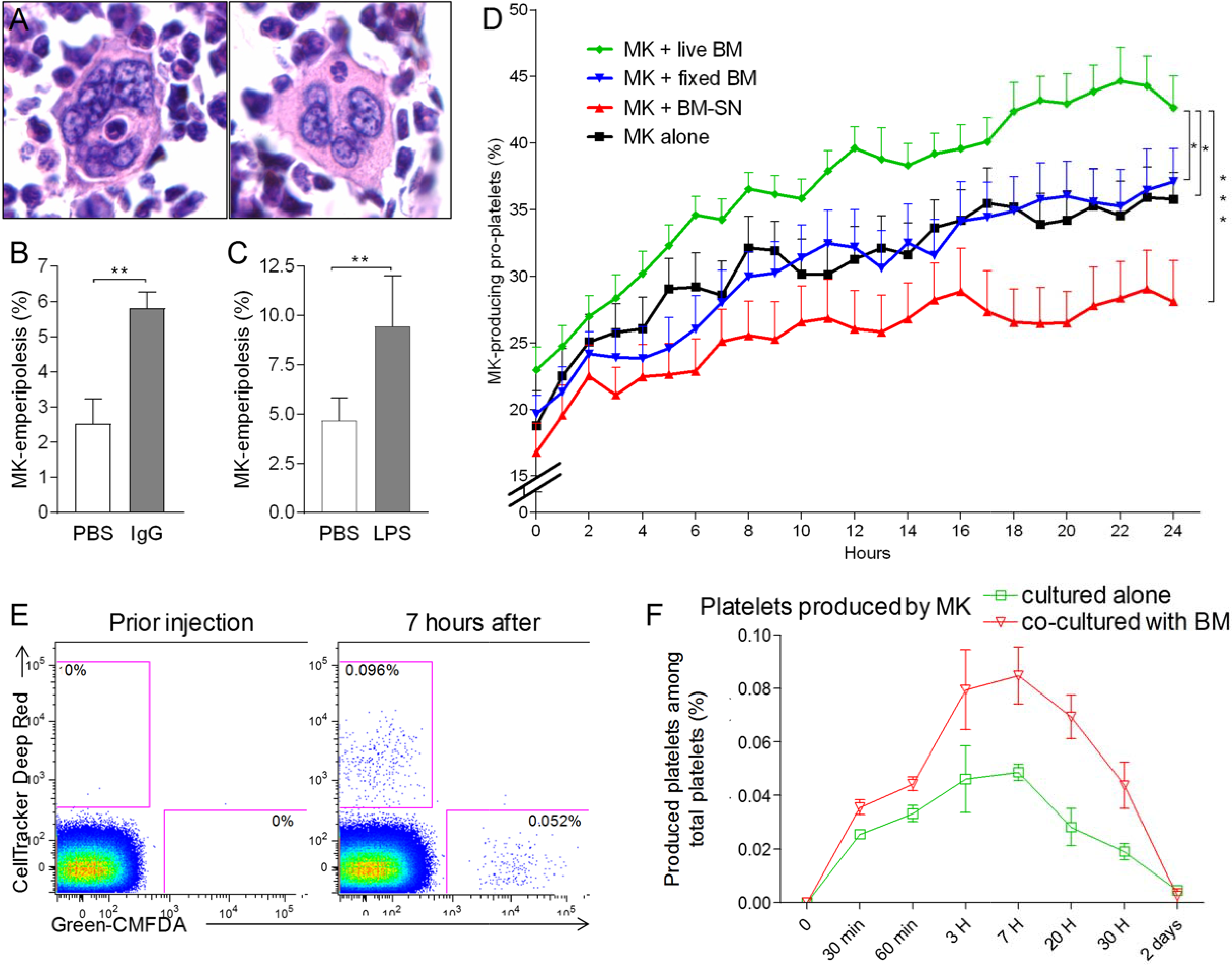
Emperipolesis contributes to platelet production. **A**. Bone sections stained with H&E showing MK containing one neutrophil in an emperiosome (left) or without evident vacuole (right). **B**. Tg-FcγR2a mice are treated with HA-IgG for 7 days. Percentage of MKs containing at least one neutrophil was calculated in bone sections, n=4-6 mice per group. C. Mice were treated with PBS or 25μg LPS i.p. After 3 days, the percentage of MKs containing at least one neutrophil is calculated in bone sections, n=4 mice per group. **D**: MKs were cultured alone (black line) or with SN from marrow cell culture (red line), fixed marrow cells (blue line) or live marrow cells (green line). MKs were then enriched and cultured in TPO medium. MKs producing pro-platelets over time were quantitated using Incucyte. None vs. live marrow: p=0.015; marrow-SN vs. live marrow: p=0.0002; fixed marrow vs. live marrow: p=0.014. Pooled from 4 independent experiments. **E-F**: MKs stained with Green-CMFDA or CellTracker Deep Red are either cocultured or not with marrow cells, mixed together 1:1 and injected i.v. (Fig S6C). After different time points, circulating platelets positive for Green-CMFDA or CellTracker Deep Red are detected by flow cytometry. **E**. Representative dot plots of green vs. red staining on CD41+ platelets. **F**. Frequency of green vs. red population of platelets over time, n=3 mice per group, representative of 3 independent experiments.

To quantitate the impact of emperipolesis on thrombocytopoiesis, we employed IncuCyte high-content live-cell microscopy (Thon et al., 2012), comparing pro-platelet generation by MKs cultured alone or together with marrow cells. These studies employed fetal liver MKs because of their superior ability to generate pro-platelets *in vitro.* To assess the role of cell-cell contact and bone marrow cell-derived soluble factors, including microparticles, we cultured MKs with marrow cell supernatant or with paraformaldehyde-fixed marrow cells. Co-culture with living marrow cells markedly enhanced pro-platelet production (Fig 6D). By contrast, MKs cultured with fixed cells or cell supernatants produced fewer pro-platelets than those cultured alone, weighing against a role for contact and soluble factors and implicating emperipolesis directly (Fig 6D).

Finally, we tested the effect of emperipolesis on platelet production *in vivo* via adoptive transfer. MKs were labeled either with Green-CMFDA or with CellTracker Deep Red, and one population or the other (varied across experiments) was cultured together with marrow cells. MKs were mixed 1:1 and engrafted IV into recipient animals for serial parallel quantitation of green and red platelets (Fig S6C). As predicted by the IncuCyte findings, MKs cultured with marrow cells were more efficient at producing platelets (Fig 6E&F), consistent with promotion of platelet production by emperipolesis.

## Discussion

Megakaryocytes anchor hemostasis via elaboration of platelets. Platelet production can occur in a cell-intrinsic manner, as for example by MKs cultured in isolation *ex vivo.* However, physiological platelet generation proceeds in a complex multicellular environment. The present studies establish a pathway through which this cellular context modulates thrombocytogenesis. During emperipolesis, neutrophils and other hematopoietic lineages penetrate into the MK cytoplasm, a process mediated actively by both host and donor. Cytoplasmic neutrophils transfer membrane and cytosolic contents to MKs and to platelets, thereby enhancing platelet production. Donor neutrophils receive membrane in turn before they exit intact (Fig S7). Thus, emperipolesis represents a previously unrecognized pathway through which neutrophils and other hematopoietic cells engage with MKs to modulate the composition and production of circulating platelets.

At a cell biological level, emperipolesis appears distinct from other cell-in-cell interactions. For example, in phagocytosis, the engulfing cell actively encompasses a passive target, typically followed destruction of the target cell when the phagosome fuses with lysosomes (Overholtzer and Brugge, 2008). In entosis, the target cell contributes actively to the engulfing event, although in most (if not all) cases destruction of the target cell is the ultimate outcome (Overholtzer and Brugge, 2008; Overholtzer et al., 2007). While the ultrastructure of these cell-in-cell phenomena remains incompletely described, target cells are thought to remain within intracellular vacuoles, rather than to enter into host cytoplasm (Fuentes et al., 2010). Membrane transfer between cells is well recognized, including via trogocytosis (Joly and Hudrisier, 2003). However, to our knowledge, entry of one cell into the cytoplasm of another to share membrane has not been previously described.

We identified β2 integrins and MK ICAM-1/ezrin as contributors to emperipolesis. These proteins also mediate another form of transcellular passage, the migration of neutrophils through the cell bodies of endothelial cells (Ley et al., 2007). Unlike emperipolesis, endothelial transcellular migration is not known to involve penetration into the host cytoplasm. It remains unknown whether other mechanisms are shared between emperipolesis and transendothelial migration, such as fusion of caveolin vesicles to create an intracellular channel for passage (Ley et al., 2007; Millan et al., 2006). Tavassoli and colleagues had previously postulated that emperipolesis could represent a pathway of neutrophil egress from the bone marrow (Dzieciol et al., 1995; Tavassoli, 1986). Transit of some neutrophils through MKs over the course of just a few minutes lends plausibility to this hypothesis. Our data do not exclude the possibility that some neutrophils pass through MKs without a cytoplasmic “detour,” thereby resembling endothelial transcellular migration even more closely.

Mechanisms of MK-emperipolesis have remained entirely obscure for more than 50 years (Larsen, 1970). Electron microscopy observations previously suggested that neutrophils are not internalized by MK but rather enter directly through the DMS (which is continuous with the cell surface (Eckly et al., 2014)) to reside within the DMS dilated cavities (Breton-Gorius and Reyes, 1976; Thiele et al., 1984). Consistent with these observation, myeloid cells are often found at the cell surface entrance of the DMS (Thiele et al., 1984), and increased emperipolesis has been reported in models with dilated and enlarged DMS, or after pharmacological modification of the DMS (Overholtzer and Brugge, 2008). However, our confocal and electron microscopy images clearly demonstrate that neutrophils can enter MKs directly, through a vacuole, ultimately taking up residence inside the MK cytoplasm. Emperipolesis is nevertheless strikingly heterogeneous. For example, emperipolesis can be observed in MKs of all sizes, with or without DMS, and can last just minutes or over an hour. MKs may enclose a single neutrophil or encompass as many as 50 neutrophils (Figure S1C). These observations strongly suggest that there may be different types of emperipolesis, involving different molecular pathways and serving distinct functions that remain to be defined.

Because the mechanistic pathways identified in our study are not specific to emperipolesis (e.g. β2 integrin binding, actin polymerization), we have so far been unable to interrupt this phenomenon with selectivity *in vivo.* To address its function, we employed a combination of approaches, focused here on platelet production. This focus is justified by the enhanced frequency of emperipolesis in diseases associated with high platelet count (essential thrombocythemia, polycythemia vera (Cashell and Buss, 1992; Vytrva et al., 2014)) or high platelet demand (gray platelet syndrome, blood loss or hemorrhagic shock (Di Buduo et al., 2016; Dzieciol et al., 1995; Larocca et al., 2015; Monteferrario et al., 2014; Sahebekhitiari and Tavassoli, 1976; Tavassoli, 1986)). Further, enhanced emperipolesis in chronic myeloproliferative disorders positively correlates with the peripheral platelet count (Thiele et al., 1984). We establish that emperipolesis accelerates platelet production both *in vitro* and *in vivo.* Whether this effect reflects enhanced access to lipid membrane or other mechanism remains to be established.

Since platelets generated through emperipolesis bear donor membrane as well as parent MK membrane, it is likely that they will be different in function. Given the role of β2 integrins in emperipolesis, it is plausible to suspect that the activated neutrophils will preferentially engage in emperipolesis, suggesting the possibility that “angry neutrophils make angry platelets”. The identity of lipids and proteins transferred from neutrophils and other cells to MKs and platelets, and the resulting changes in cell function, will be important topics for future study.

The impact of emperipolesis is unlikely to be restricted to MKs and platelets. Our videomicroscopy data confirm that exiting neutrophils can carry MK membrane with them (Fig S4D), potentially translating into altered function. These effects are more difficult to study *in vitro* because of the capacity we identified for MKs to transfer membrane to nearby cells via MK microparticles (Fig S4H). Our prior work had identified MK microparticles as potent pro-inflammatory vectors, potentially mediating delivery of IL-1 in systemic inflammatory disease (Cunin et al., 2017). The present findings thus extend the understanding of MK microparticles as signaling vectors. Further, by establishing transfer of membrane not only from donor cell to MK but also reciprocally, followed by release of viable cells back into the intercellular milieu, these studies identify emperipolesis as a mechanism through which MKs may be able to “groom” neutrophils and other immune lineages.

We are unable at present to define conclusively the proportion of circulating platelets that bear neutrophil membrane. The mT/mG chimera experiments will not accurately reflect this fraction, because only one of 4 possible donor-host pairs yields detectable platelets (mT/mG neutrophil➔WT MK) and because mT/mG fluorescence is weak, such that transfer events will likely often be invisible. We note that many platelets released by MKs co-cultured with membrane-labeled marrow donors expressed membrane label (Fig 5). Given the speed with which cells enter and exit MKs, a snapshot prevalence of 6% is compatible with the possibility that many or even most MKs, and many neutrophils, experience emperipolesis over time, perhaps repeatedly. If this is the case, and labeling experiments accurately reflect the efficiency of membrane transfer, then emperipolesis-derived membrane could be common in circulating platelets. Alternately, if transfer were inefficient, and/or only a subset of MKs engaged in emperipolesis, then emperipolesis-modulated platelets could represent simply a small (but potentially still functionally important) subset of the circulating pool.

We recognize other limitations to these studies. The function of emperipolesis-derived platelets remains to be established. The mechanisms through which neutrophils escape emperisomes to enter the cytoplasm, home intracellularly to the DMS, and then egress without violating MK outer membrane integrity remain to be defined. Cells deficient in β2 integrins retain the capacity to enter MKs, albeit with reduced efficiency, revealing that other ligand/receptor pairs can mediate entry. The signals driving enhanced emperipolesis in the setting of experimental stress, during hemorrhagic shock, and in aberrant marrow environments such as in hematopoietic malignancies, remain to be established. Despite these limitations, the current studies identify emperipolesis as a novel cell-in-cell interaction that mediates reciprocal transfer of membrane and other cellular components, defining thereby a new mechanism of interchange between immune and hematopoietic systems.

## Acknowledgements

### Funding

This work was supported by the Arthritis National Research Foundation (to PC); NIH awards 5F32HL118865, K01DK111515, and American Heart Association 16SDG29090007 (to KRM); NIH award R01H168130 (to JEI); and NIH awards R21AR062328, R01AR065538, and P30AR070253, the Cogan Family Foundation, and the Fundación Bechara (to PAN). KRM is an American Society of Hematology Scholar.

### Data and materials availability

All data is available in the main text or the supplementary materials

## Author contribution

P.C. conceived the project, performed experiments, analyzed data, and wrote the manuscript. R.B., K.R.M., M.M.B., N.N.M., A.M., and E.B. performed experiments. K.R.M, M.M.B., P.Y.L., A.G., A.W., M.S.V., E.B. and J.E.I. discussed the results, commented on the manuscript and contributed important intellectual insights. P.A.N. conceived the project, supervised the work, analyzed data, and wrote the manuscript.

## Supplemental Materials

### Materials and Methods

#### Mice

C57Bl/6, CD45.1 B6 mice, CD18^-/-^ mice, mT/mG mice and Tg-FcγRIIA mice were purchased from The Jackson Laboratory. LyzM-GFP mice transgenic for FcγRIIA (Tg-FcγRIIA) mice were backcrossed 10 times in the C57BL/6J background. Unless stated, all experiments employed male mice aged 8-12 weeks. All procedures were approved by the local animal care committee.

#### Antibodies

Anti-CD61 (2C9.G2), -CD42d (1C2), -CD11a (M17/4), -CD11b (M1/70), CD62P (RMP-1), CD31 (MEC13.3, all from Biolegend), -Ly6G (1A8, BioXCell), -GPVI (Jaq1, Emfret), and anti-CD18 (clone GAME-46, BD biosciences) were used in blocking experiments. Antibodies used for flow cytometry and microscopy staining were anti mouse-CD41 (MWReg30), -ICAM (YN1/1.7.4), -CD18 (M18/2), -Ly6G (1A8), -CD31 (MEC13.3), CD144 (BV13), -Tubulin (10D8 all from Biolegend), -Ezrin and -phospho-Ezrin (rabbit polyclonal, Cell Signaling), and anti-human-CD41 (HIP8), -CD15 (W6D3), CD66b (G10F5), and -ICAM-1 (HA58). Fluorescent-conjugated secondary antibodies were purchased from Jackson Immunoresearch.

#### Chemicals and reagents

Latrunculin A and Cytochalasin D were purchased from Cayman Chemical. Ezrin inhibitor NSC668394 was from Calbiochem. Lipids cells strainers PKH67 and PKH26 were from Sigma. Protein strainers Green CMFDA, CellTracker Deep Red and Cell Trace Violet were from Molecular probes. Surface protein biotinylation kit was purchased from Pierce.

#### Cell generation

Murine MK generation: Bone marrow cells were cultured in complete RPMI medium supplemented with 1% supernatant from the TPO-producing fibroblast cell line GP122 (hereafter called TPO medium) (Villeval et al., 1997). After 4-5 days, MKs were separated from marrow cells using a two-step albumin gradient as described (Schulze, 2016; Shivdasani and Schulze, 2005). Fetal liver-derived MKs were generated as described (Machlus et al., 2017). Human MK generation: mobilized peripheral blood or BM CD34+ stem/progenitor cells were purchased from AllCells. 1×10^5^ cells were cultured in StemSpan medium supplemented with a MK expansion supplement, both from StemCell for 12-14 days, as described (Liu et al., 2011). Human neutrophils were obtained from blood from healthy donors. Neutrophil were enriched using a dextran density gradient sedimentation as described (Cunin et al., 2016) Contaminating red blood cells were lysed by hypo-osmotic shock.

#### Emperipolesis assay

2×10^4^ murine MKs with 2×10^6^ murine marrow cells in TPO medium, or 2×10^4^ human MKs with 2×10^6^ human neutrophil in StemSpan medium were co-cultured overnight in P96 round bottom wells.

#### Confocal microscopy

Cells were fixed in PFA 2% for 30 min at RT. After washing, cells were suspended in PBS supplemented with 0.1% saponin and 3% FCS (permeabilization buffer) and incubated 2 hours at RT or overnight at 4C with 10μg/ml primary antibodies. After washings in permeabilization buffer, secondary antibodies diluted 1:200 were added for 1 hour. When indicated, Phalloidin (Molecular Probe), Draq5 (eBioscience) or Hoechst (ThermoFisher) were added for the 15 last minutes, prior washing with PBS and cytospin. Cells were mounted on slides using FluorSave mounting medium (Calbiochem). Microscopy was performed using a Nikon C1 Plus Confocal Laser Scanner confocal or a Zeiss LSM 710 or 800 Multiphoton Laser scanner confocal microscope.

#### Spinning disk confocal microscopy

MKs stained with PKH26 and marrow cells stained with PKH67 were co-cultured in P96 round bottom wells for at least 1 hour prior spinning disk assay. Cells were then resuspended in TPO medium without red phenol and supplemented with Draq5, and cultured in a micro insert 4 wells dish for time lapse imaging. Movies were obtained using a YokogawaCSU-X1 or an Olympus DSU inverted spinning disk confocal microscope. Images were acquired every 4 minutes on 12 different Z-stacks, 1μm per stack. Movies were analyzed using EZ element software or Volocity software.

#### Imaging of whole-mount bone marrow

Whole-mount-tissue preparation, immunofluorescence staining and imaging of the bone marrows were performed as described previously (Kunisaki et al., 2013). Briefly, mice were intravenously injected with AF647-labelled anti-CD31 and anti-CD144 and perfused with PBS and 4% PFA15 min after. Femurs and tibias were harvested, PFA-fixed, frozen in OCT, and shaved on a cryostat to expose the marrow. Bones were incubated in PBS containing 10%FCS and 0.5% Triton X-100, with AF594 anti-Ly6G, AF488 anti-CD41 and Hoechst for 2 days. Images are acquired using a Zeiss LSM 800 Multiphoton Laser scanner confocal microscope and reconstructed in 3D with Imaris software.

#### Electron microscopy

Cells were fixed using 2% PFA and 0.1% glutaraldehyde for 1 hour RT. After washings, cells were incubated with 0.1% OsO4 for 30 minutes prior to sectioning. 50nm sections were observed with a JEOL 1200EX electron microscope.

#### In vitro pro-platelet production

MKs were co-cultured overnight without or with PKH67-stained marrow cells. Marrow cells were separated from MKs using BSA-gradient sedimentation, and cells were transferred to a P24 well plate and imaged using the IncuCyte HD system (Essen BioScience). Frames were captured every hour. Rates and extent of proplatelet production were measured in ImageJ software using investigator-coded software (Thon et al., 2012).

#### In vivo platelet production

2×10^5^ MKs, previously stained with Green CMFDA or CellTracker Deep Red (Molecular probes) and co-cultured or not with marrow cells, were injected i.v. in 200μl PBS. Blood was harvested by tail vein sampling at indicated time points using heparinized capillary tubes. 1μl was blood is diluted in 500μl PBS in the presence of an anti-CD41 antibody. Presence of green CMFDA or CellTracker Deep Red on circulating CD41+ platelets was evaluated by flow cytometry.

#### Emperipolesis in marrow sections

Bones were fixed in PFA 4% for 2 days prior to decalcification in Kristensen solution for 2 days prior to paraffin-embedding. Percentages of marrow MKs containing at least one neutrophil were determined on 6μm paraffin-embedded sections stained with H&E.

#### Platelet overproduction models

*LPS treatment:* WT C57Bl/6J mice were treated i.p. with PBS or 25μg LPS in 200μl PBS. Bones were harvested 3 days later. *FcγRIIA transgenic mice:* Tg-FcγRIIA mice were treated i.v. with PBS or 500μg heat-agglutinated IgG. Bones were harvested 7 days later. *TPO administration:* WT mice were treated daily with 0.5μg rmTPO (Peprotech) or PBS i.v. for 3 days (Kirito et al., 2002). Bones were harvested 7 days later. *Immune thrombocytopenia:* WT mice were treated i.v. with 5μg anti-CD41 (clone MWReg30) or isotype control (Rat IgG1 clone RTK2071) (Hitchcock et al., 2008). Bones were harvested 2 days later. Circulating CD41+ platelet were quantified by flow cytometry using 1μm counting beads (Polysciences, Inc).

*Statistics:* statistical significance in emperipolesis between 2 conditions was determined using the Chi-square test, with at least 150 MKs counted per sample. To compare emperipolesis in 2 groups of mice we used the Mann-Whitney test. For other experiments, we used the Wilcoxon matched pairs test. All statistical analysis were done using Prism software, * p<0.05, ** p<0.01 *** p<0.001.

## Supplemental Figures

**Figure S1.**
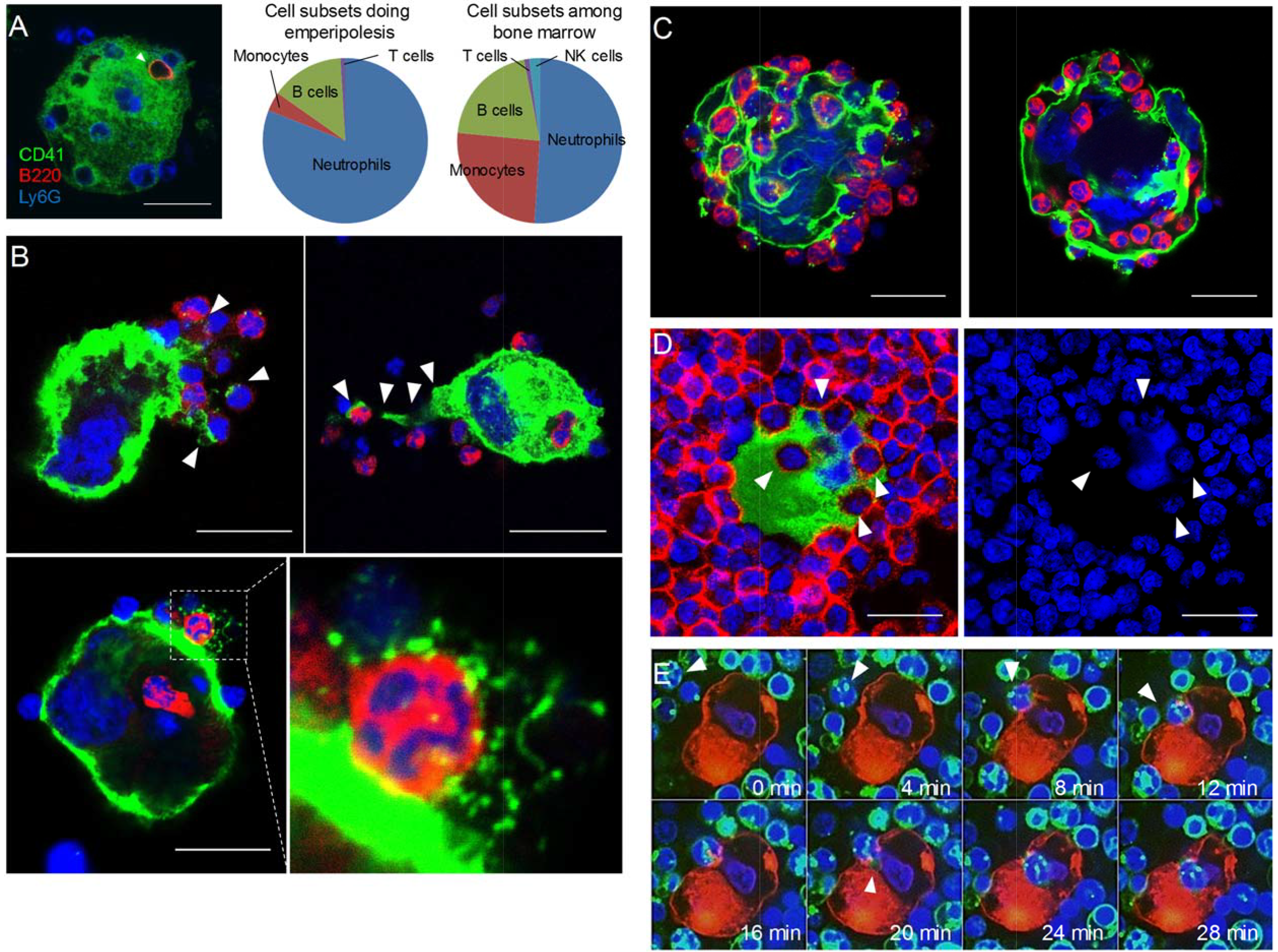
Visualization of murine and human emperipolesis by confocal microscopy. **A**. Murine MK and marrow cells were co-cultured and stained with anti-CD41 (green), anti-Ly6G (blue), and either anti-CD3ε, -B220 (red), -CD115, -NK1.1, or anti-Kit (red). Left: confocal images showing a B cell and neutrophils inside MK cytoplasm. Middle: marrow cell subsets undergoing emperipolesis. Right: cell subset percentages after overnight incubation in culture medium. **B-C**. After co-culture, cells were stained with anti-CD41 (green), anti-CD18 or –Ly6G (red) and Draq5 (DNA, blue). **B**. Examples of MK tethers surrounding neutrophils or other marrow cells (see also Supplemental movie 2). **C**. MKs containing more than 20 neutrophils. **D**. Cells from human bone marrow aspirate were stained with anti-CD41 (green), anti-CD66b (red), and Draq5 (blue). The image on the right shows DNA staining only, allowing to visualize neutrophils near the MK nucleus (can also be observed in Fig 1B, and in mouse in Supplemental movie 1). **E**. Murine MKs stained with PKH26 (red) and marrow cells stained with PKH67 (green) were co-cultured in the presence of DraqV for time-lapse spinning disk microscopy. (see also Supplemental movie 3)

**Figure S2:**
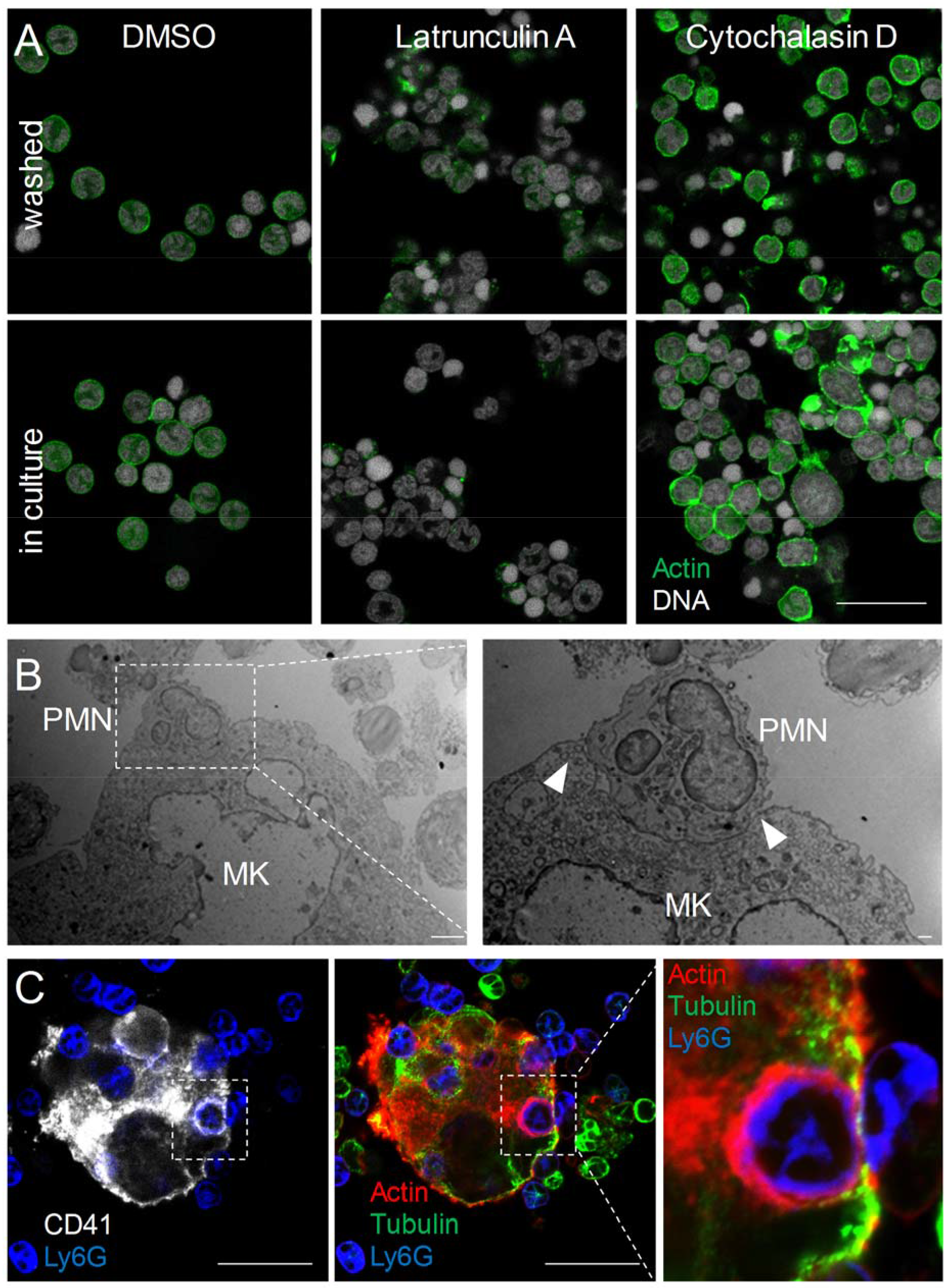
*In vitro* modeling of emperipolesis reveals a rapid multi-stage process. **A**. Marrow cells were treated with 1μM latrunculin A, cytochalasin D, or DMSO. After 2 hours, cells were either washed thoroughly to remove inhibitors and cultured overnight (upper photos) or left in culture in the presence of the inhibitors (lower photos). Images show a strong disruption of F-actin in cells treated with latrunculin A, and bright actin aggregates distributed irregularly within the cells with cytochalasin D, as reported (Khajah et al., 2015; Rubtsova et al., 1998). **B**. Electron microscopy images showing the formation of a cup on MK surface. Scale bars: 2μm (left) and 500nm (right). **C**. After co-culture, cells were stained with anti-CD41 (white), anti-Ly6G (blue), anti-tubulin (green). Actin was visualized with phalloidin (red). **A&C**. Scale bars represent 20μm.

**Figure S3:**
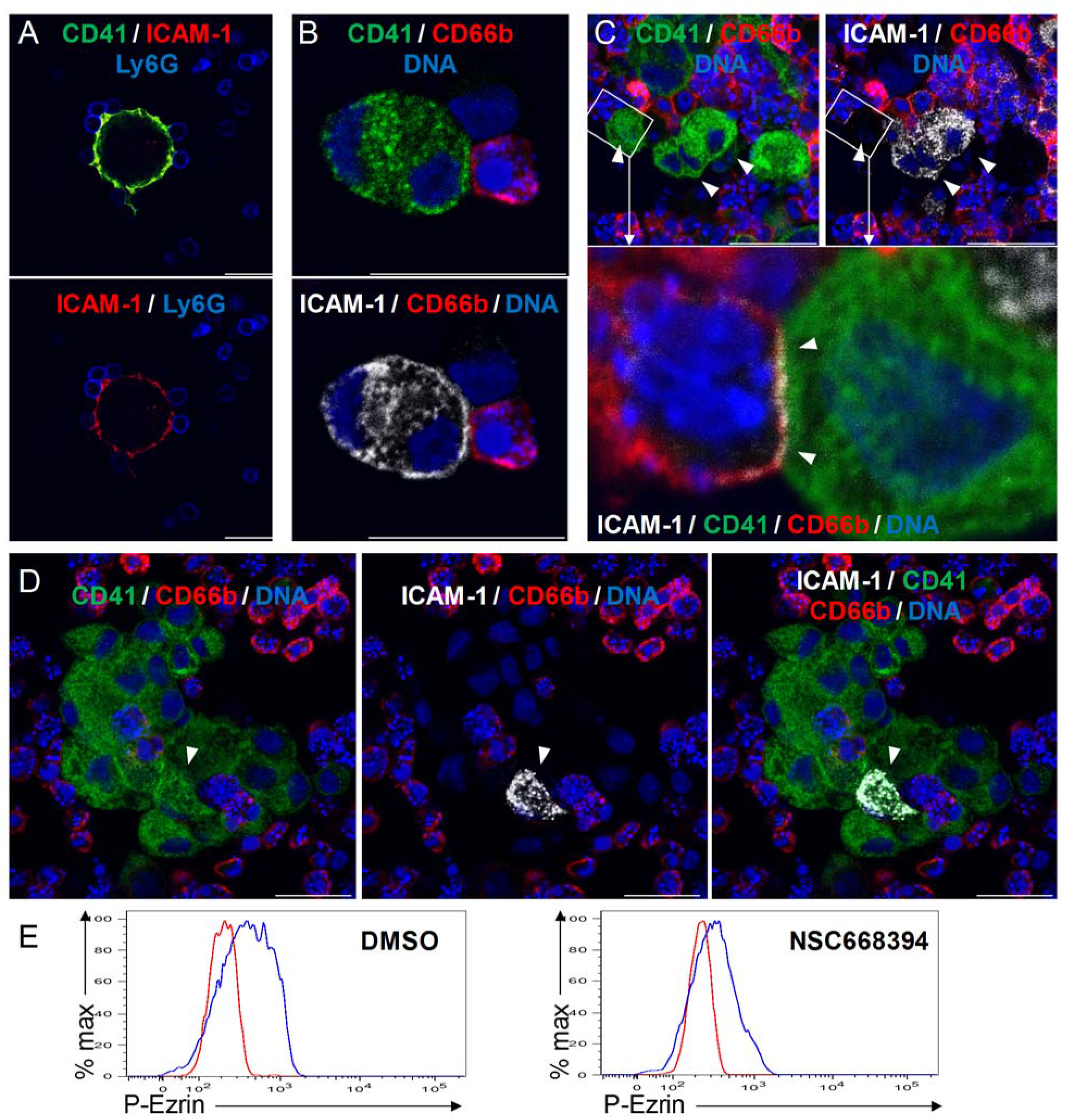
Emperipolesis is mediated by active actin cytoskeleton rearrangement and in part through β2-integrin/ICAM-1/ezrin. **A**. MKs generated from murine marrow were co-cultured with marrow cells. **B-D**. MKs generated from human CD34+ mobilized cells were co-cultured with peripheral blood neutrophils. **A-D**. After fixation, cells were stained with anti-CD41, anti-Ly6G or -CD66b, and anti-ICAM-1. DNA was visualized with Hoechst. ICAM-1 was observed on a fraction of murine and human MKs (~5%, arrowheads), principally those engaged with in cell-cell contact with marrow cells. Scale bars represent 20μm. **E**. Marrow cells were treated overnight with DMSO (left) or 1μM of the ezrin inhibitor NSC668394 (right), PFA-fixed, permeabilized and incubated with an anti-phosphorylated ezrin antibody or a rabbit isotype control overnight at 4C. After washing, cells were incubated with a AF488-conjugated anti-rabbit Ab. Histograms show phospho-ezrin MFI (blue) vs. isotype control MFI (red).

**Figure S4:**
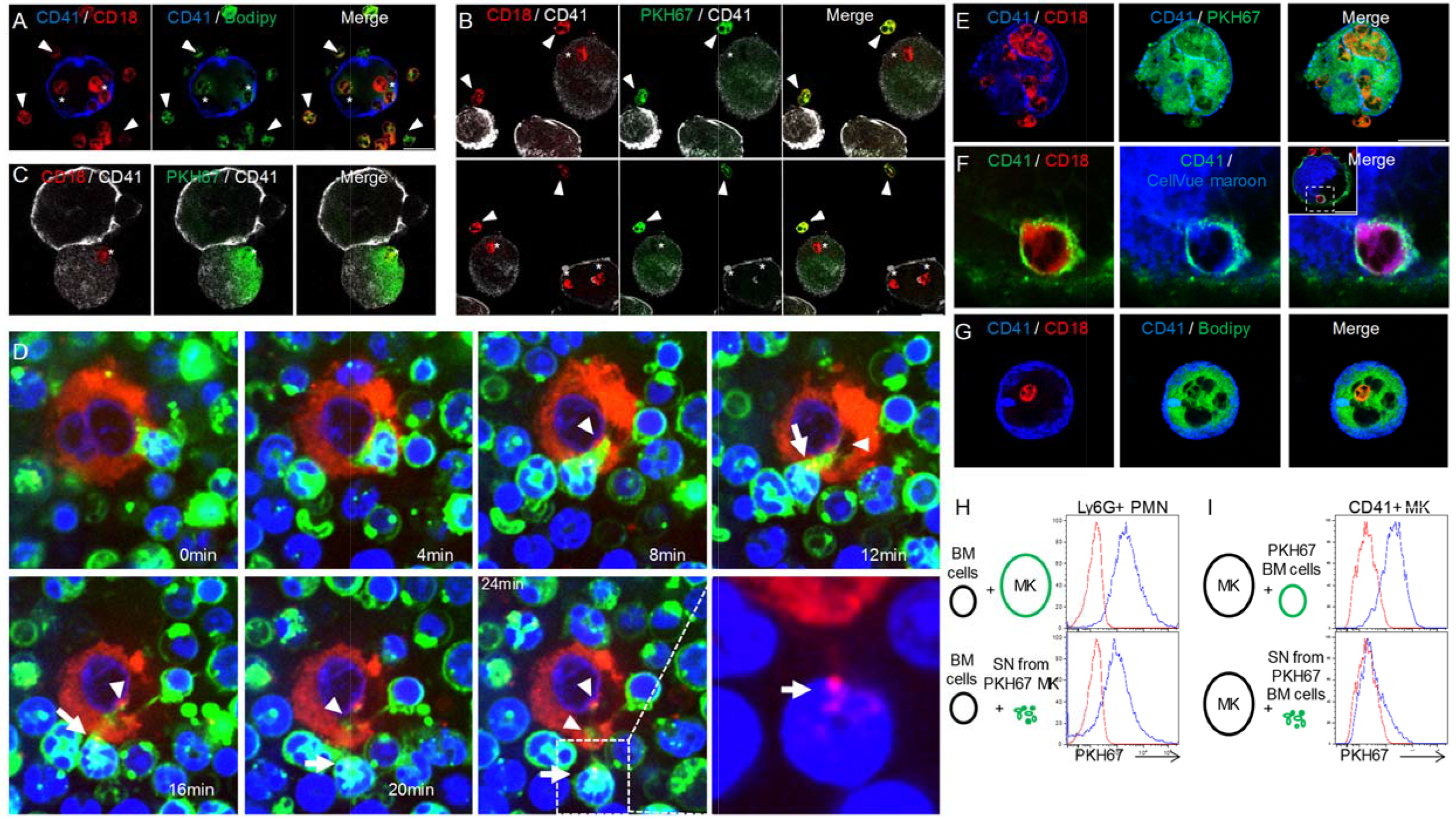
Emperipolesis mediates membrane transfer from neutrophil to megakaryocyte. Marrow cells were stained with Bodipy (**A**) or PKH67 (**B&C**) prior to emperipolesis assay. Confocal microscopy shows a loss of lipid stainer fluorescence in neutrophils engaged emperipolesis (asterisk). Arrowheads show cells with intact PKH67 fluorescence. **C**. Enhanced green settings reveal a presence of PKH67 into MK, with an appearance of diffusion into cytoplasm. **D**. Murine MKs stained with PKH26 (red) and marrow cells stained with PKH67 (green) are co-cultured in the presence of DraqV. Time-lapse spinning disk microscopy revealed a rapid passage of a green neutrophil inside MK, leaving green membrane behind (arrowheads). MK red membrane is observed on neutrophil after emperipolesis (arrows and lower right photo without green staining). (see also Supplemental movie 4). **E-G**. MKs were stained with (**E**) PKH67, (**F**) CellVue Maroon, or (**G**) Bodipy prior to co-culture with marrow cells. Confocal microscopy shows co-localization between lipid stain and neutrophil membrane. **H**. Marrow cells were cultured with PKH67+ MKs (upper panel) or supernatant (SN) from PKH67+ MK culture (lower panel). Histograms show PKH67 MFI on Ly6G+ neutrophils after culture with unstained MK/SN (red histograms) or PKH67+ MK/SN (blue histograms). **I**. MKs were cultured with PKH67+ marrow cells (upper panel) or supernatant from PKH67+ marrow cell culture (lower panel). Histograms show PKH67 MFI on CD41+ MK after culture with unstained marrow cells/SN (red histograms) or PKH67+ marrow cells/SN (blue histograms).

**Figure S5:**
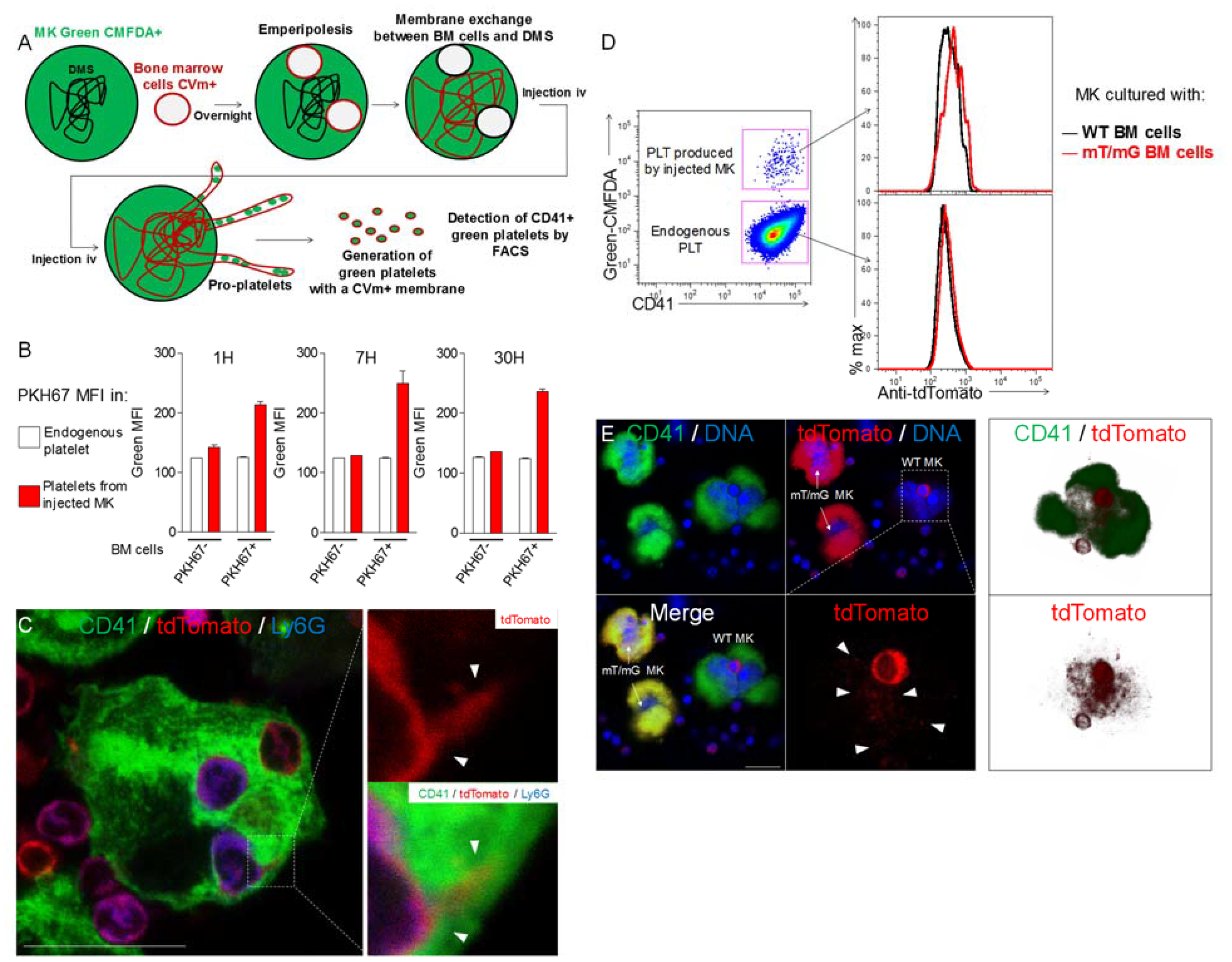
Neutrophil membranes transferred in emperipolesis emerge on circulating platelets. **A**. Experimental design for platelet production *in vivo.* MKs are stained with green-CMFDA to monitor platelet production. CMFDA+ MKs are co-cultured with marrow cells stained with CellVue maroon (far-red lipid stain). If membrane transfer occurs from neutrophil to MK DMS, MKs injected i.v. will produce green platelets bearing far-red (CellVue maroon+) membrane. Tail vein sampling was performed at different time points in the presence of heparin, and fluorescence on green platelets was studied by flow cytometry. **B**. Same as in Figure 5A&B, but MKs are stained with CellTracker Deep Red instead of green-CMFDA, and marrow cell membrane is stained with PKH67 instead of CellVue maroon. Presence of PKH67 on platelet surface is studied by flow cytometry. **C**. MKs were co-cultured overnight with marrow cells from mT/mG mice. Cells were stained for CD41 (green), Ly6G (blue) and tdTomato (red). Images show the appearance of tdTomato+ Ly6G-signal within MKs. **D**. MKs stained with green-CMFDA and co-cultured with WT or mT/mG BM cells were injected i.v. After 1 hour, circulating platelets are fixed and stained with an anti-tdTomato antibody. Right: Green-CMFDA stain distinguishes platelets produced by injected MKs from endogenous platelets. Left: tdTomato MFI on platelets in mice injected with MKs co-cultured with WT cells (black histogram) or mT/mG cells (red histograms). Representative of 2 experiments. **E**. Bone marrow from chimeric mice was flushed directly into PFA, and cells were stained with CD41 (green) and tdTomato (red). DNA is visualized with Hoechst. Right: 2 mT/mG MKs and 1 WT MK containing a mT/mG marrow cell are shown. tdTomato signal is detected around the mT/mG cell (lower right photo). Right: 3D reconstitution of tdTomato signal inside the WT MK.

**Figure S6:**
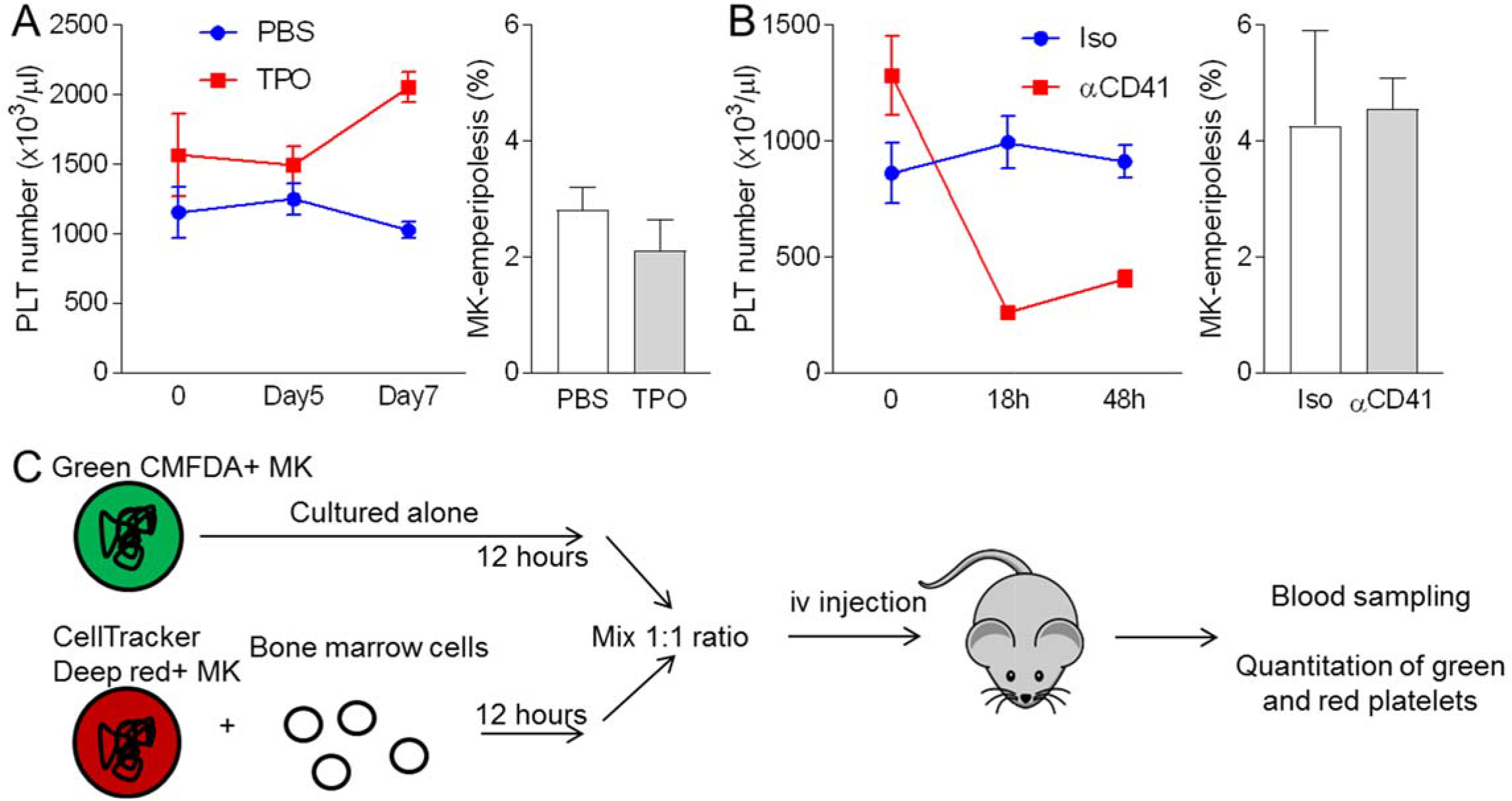
Emperipolesis contributes to platelet production. **A**. Mice were injected daily with PBS or 0.5μg rmTPO i.v. for 3 days, and bones were harvested after 7 days. **B**. Mice were injected with 5μg anti-CD41 or isotype control i.p. and bones were harvested after 2 days. **A-B**. Left: circulating platelet count over time after TPO (**A**) or anti-CD41 (**B**) treatment. Right: percentages of MKs containing neutrophil in bone sections stained with H&E. n=3 females (**A**) and 3 males (**B**) per group, at least 400 MKs per group were counted. **C**. Experimental procedure for MK engraftment (Fig 6E&F). Briefly, MKs stained with Green-CMFDA or CellTracker Deep Red were cultured alone or with marrow cells, respectively. After 12 hours incubation, MKs were counted and mixed at a 1:1 ratio. At least 200,000 MKs per recipient were injected i.v.

**Figure S7:**
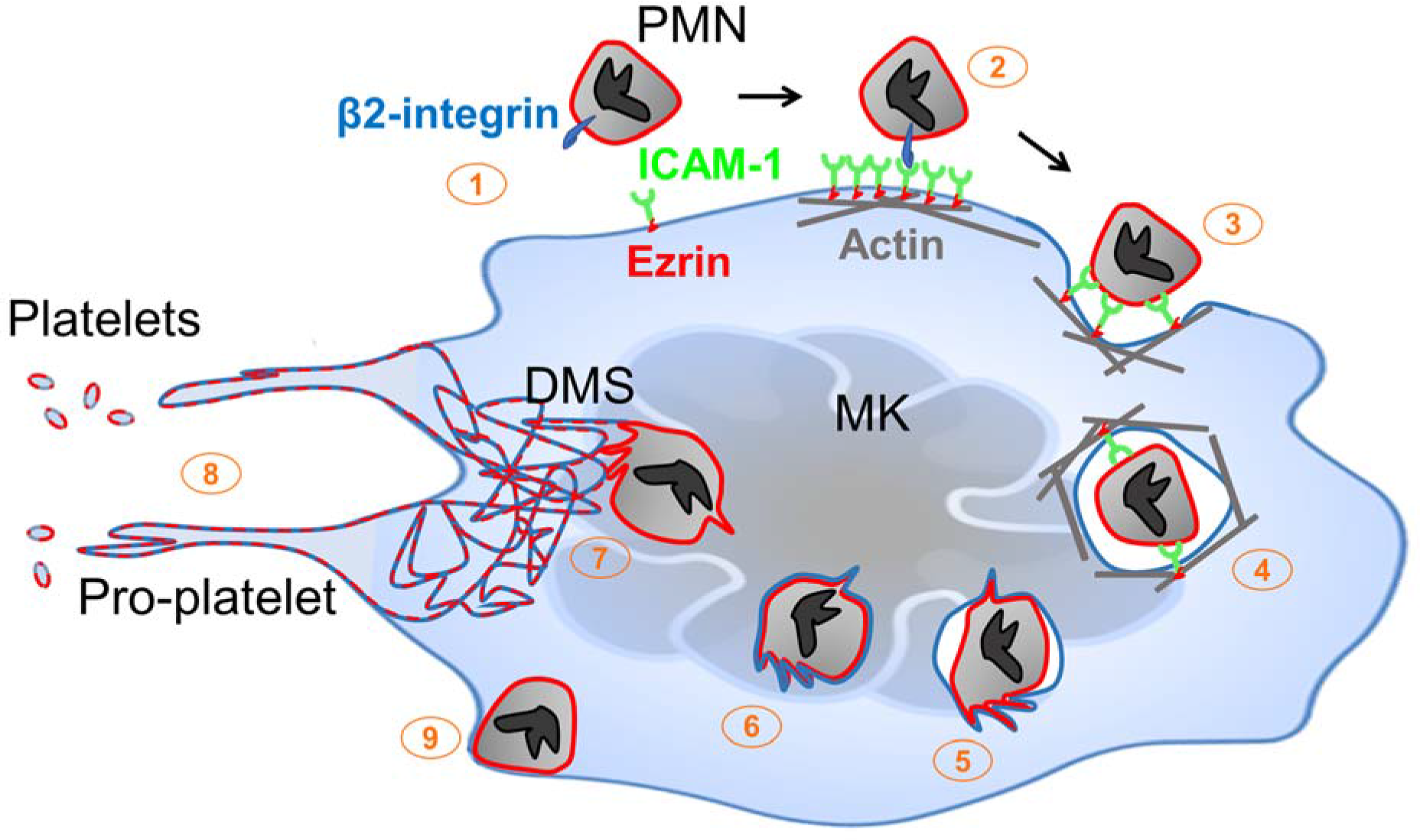
Neutrophil transit through megakaryocytes in emperipolesis mediates membrane transfer and enhanced platelet production – proposed model. **1**. Neutrophils expressing β2 integrin and MKs expressing ICAM-1 are positioned to engage, including via MK tethers. **2**. ICAM-1 ligation by neutrophil β2-integrins induces ICAM-1/ezrin pair translocation and clustering, as in transendothelial migration (Ley et al., 2007). **3**. Formation of a trans-megakaryocyte cup in an actin-dependent manner. **4**. Neutrophil entry into MK vacuole (“emperisome”). **5-6**. Neutrophil and emperisome membranes become closely apposed. Neutrophil and emperisome membranes may merge at some locations. Emperiosome membrane disappears, enabling neutrophil entry into the MK cytoplasm. **7**. Neutrophil translocates to DMS where membrane fusion occurs, transferring neutrophil membrane and protein. **8**. Production of platelets containing neutrophil membrane. **9**. Egress of viable neutrophil from MK.

## Supplemental Videos

**Movie 1: Emperipolesis within murine bone marrow.** Three-dimensional reconstitution of murine marrow, showing MKs (green) neutrophils (red), bone marrow sinusoids (white), and DNA (blue). Green, red or blue fluorescence are removed occasionally to visualize neutrophils inside MK or MK tethers. The 3 animations correspond to the 3 images shown in Figure 1A.

**Movie 2: Neutrophil enters megakaryocyte - formation of a trans-megakaryocyte cup.** MKs stained with PKH67 (green) were co-cultured with marrow cells stained with PKH26 (red) in the presence of Draq5 (DNA, blue). Video shows the formation of a transcellular cup on the MK surface allowing neutrophil entry.

**Movie 3: Megakaryocyte tethers and neutrophil entry into a CD41+ vacuole.** MKs stained with anti-CD41 (green) were co-cultured with marrow cells from mT/mG mice (red) in the presence of Draq5 (DNA, blue). Video shows a neutrophil on the MK surface, attached by MK tethers, followed by a rapid entry through a CD41+ membrane. A few minutes after its entry, the neutrophil exits at the bottom of the field of view.

**Movie 4: Neutrophil exits megakaryocyte.** MKs stained with anti-CD41 (green) were co-cultured with marrow cells from mT/mG mice (red) in the presence of Draq5 (DNA, blue). A neutrophil is present inside megakaryocyte, but without interposed CD41+ membrane. Neutrophil exits MK and remains attached on its surface. Video obtained with a laser scanning confocal microscope.

**Movie 5: Transfer of membrane during rapid emperipolesis.** MKs stained with PKH67 (green) were co-cultured with marrow cells stained with PKH26 (red) in the presence of Draq5 (DNA, blue). Video shows a neutrophil entering and rapidly transiting through a MK, leaving green membrane behind. Green or red fluorescence is removed at some time points to visualize bi-directional membrane transfer.

**Movie 6: Transfer of membrane during long-lasting emperipolesis.** MKs stained with PKH67 (green) were co-cultured with marrow cells stained with PKH26 (red) in the presence of Draq5 (DNA, blue). Video shows a neutrophil residing within an MK. Green or red fluorescence is removed at some time points to visualize bi-directional membrane transfer.

